# Evolution of the highly repetitive PEVK region of titin across mammals

**DOI:** 10.1101/411959

**Authors:** Kathleen Muenzen, Jenna Monroy, Findley R. Finseth

## Abstract

The protein titin plays a key role in vertebrate muscle where it acts like a giant molecular spring. Despite its importance and conservation over vertebrate evolution, a lack of high quality annotations in non-model species makes comparative evolutionary studies of titin challenging. The PEVK region of titin—named for its high proportion of Pro-Glu-Val-Lys amino acids—is particularly difficult to annotate due to its abundance of alternatively spliced isoforms and short, highly repetitive exons. To understand PEVK evolution across mammals, we first developed a bioinformatics tool, PEVK_Finder, to annotate PEVK exons from genomic sequences of titin and then applied it to a diverse set of mammals. PEVK_Finder consistently outperforms standard annotation tools across a broad range of conditions and improves annotations of the PEVK region in non-model mammalian species. We find that the PEVK region can be divided into two subregions (PEVK-N, PEVK-C) with distinct patterns of evolutionary constraint and divergence. The bipartite nature of the PEVK region has implications for titin diversification. In the PEVK-N region, certain exons are conserved and may be essential, but natural selection also acts on particular codons. This region is also rich in glutamate and may contribute to actin binding. In the PEVK-C, exons are more homogenous and length variation of the PEVK region may provide the raw material for evolutionary adaptation in titin function. Taken together, we find that the very complexity that makes titin a challenge for annotation tools may also promote evolutionary adaptation.

## INTRODUCTION

One goal of modern biology is to connect the underlying molecular structure of a protein with physiological function. Studies that compare protein sequences in an evolutionary context can illuminate the regions of proteins that are most essential, under functional constraint, or responding to natural selection (e.g., Perutz 1983; Hughes and Nei 1989; Kreitman and Akashi 1995; Galindo et al., 2003; Zhao et al., 2009; Finseth et al., 2015). With the advent of next generation sequencing, the focus of such studies has often moved beyond single gene analyses to comparing entire genomes or transcriptomes (e.g., Finseth et al., 2014; Karlson et al., 2014; Zhang et al. 2014; Pervouchine et al., 2015; Chikina et al, 2016; Partha et al., 2017). Yet, genes that are large, repetitive, and/or poorly annotated can be left behind by this approach. One such example is titin (TTN), a giant filamentous protein expressed in the muscles of all bilaterian metazoans (Steinmetz et al., 2012) that plays a key role in muscle elasticity (Linke, 2018).

At nearly 4,000 kDa and greater than 1 micron in length, titin (also known as connectin) is among the largest proteins found in vertebrates (Bang et al., 2001). Titin plays a key role in vertebrate striated muscle, where it acts as a giant molecular spring responsible for passive and active muscle elasticity (reviewed in Linke, 2018). Titin spans an entire half sarcomere from z-disk to m-line and its I-band region is composed of three domains: the proximal tandem Ig segment; the unique N2A (skeletal muscle), N2B (cardiac muscle), or N2BA (cardiac muscle) sequence; and the PEVK region (Linke et al., 1996; Gregorio et al., 1999). Together, these act as serially linked springs (Labeit and Kolmerer, 1995).

The general role titin plays in sarcomere structure is conserved among vertebrates; however, the elastic and physiological properties of vertebrate muscle vary dramatically across species and tissues. Titin, and the I-band in particular, are at least partially responsible for some of these functional differences (Freiburg et al., 2000, Prado et al., 2005, Trombitas et al., 2000) and likely contributes to variation in muscle physiology among tetrapods (Manteca et al., 2017). For example, cysteine residues in the I-band of titin may be responsible for mechanochemical evolution of vertebrate titin (Manteca et al., 2017). Likewise, variation in the length and structure of alternatively spliced titin isoforms can affect passive and active stiffness of muscles within a single species (Freiburg et al., 2000; Linke et al., 1998, Prado et al., 2005, Powers et al., 2016, Monroy et al., 2017).

Despite its central role in muscle physiology and evolution, the TTN gene is often poorly or partially annotated in non-model species. One issue is the sheer size of TTN; it is composed of more than 100 kbp with a protein length of > 30,000 amino acids and the gene is more than 60 times the length of the average eukaryotic gene (Bang et al., 2001, Granzier et al., 2007). TTN also encodes 364 exons with lengths that range from just a few base pairs to greater than 17,000 base pairs, and theoretically can produce more than one million splice variants (Bang et al., 2001, Guo et al., 2010). The abundance of alternatively spliced isoforms means that numerous tissues, samples, and individuals are required for complete cDNA-based or RNA-based annotations.

The PEVK region of TTN, an important determinant of titin and muscle elasticity, also presents a problem for most annotation tools. The PEVK region contains over 100 short, repetitive exons consisting of ~ 70% proline (P), glutamate (E), valine (V), and lysine (K) residues. These features mean that PEVK exons are often missed by automated annotation tools and previous studies have relied upon manual annotation of this region (e.g., Freiburg et al. 2000; Granzier et al., 2007). Nevertheless, the PEVK region is potentially an important target of selection over evolutionary time; it is evolutionarily labile, varies in length, exon structure, and amino acid content across some vertebrates, and may contribute to evolutionary adaptations in myofibril and whole muscle stiffness (Witt et al., 1998; Greaser et al., 2002; Granzier et al., 2007). The PEVK region also appears to have a hierarchical structure, with greater sequence divergence in the N-terminal among vertebrates than the C-terminal (Witt et al., 1998). With the wealth of genomic data now available, more detailed analyses of the PEVK region across vertebrates are possible and key to understanding its role in muscle evolution.

Here, we characterize the genomic structure of the PEVK region of TTN to gain insight into the structure-function relationships of I-band titin and its evolution across mammals. We first developed a custom tool, PEVK_Finder, to annotate exons within the PEVK region across diverse mammalian species. We then compared exon structure and sequence content both within one individual’s PEVK region and across species. Finally, we performed evolutionary analyses to examine the nature of selection acting on the PEVK region of TTN.

## MATERIALS AND METHODS

### PEVK_Finder development and optimization

We developed a tool called PEVK_Finder to annotate the PEVK region of TTN across vertebrates (Table S1). PEVK_Finder annotates exons according to three criteria: 1) minimum exon length (12), 2) minimum PEVK ratio per exon (0.54), and 3) sliding window length (10; determination of optimal parameter values described below). For a given species, PEVK_Finder translates the complete TTN sequence into three forward reading frames and splits these into separate FASTA files. PEVK_Finder then utilizes a sliding window approach to identify windows with a PEVK ratio above a minimum threshold. Note that the PEVK_Finder algorithm also incorporates the amino acid alanine (A) in its PEVK ratio calculations, based on previously observed PPAK motifs in titin (Greaser et al., 2002). However, all ratios reported in the text and figures of this paper will be referred to simply as “PEVK ratio.” All windows that meet the PEVK ratio requirement are stored, and overlapping windows are combined into discrete sequences with unique start and end coordinates. To determine exonic boundaries, PEVK_Finder searches for paired donor and acceptor splice sites within each sequence using canonical mammalian nucleotide splicing patterns (Burset et al, 2000). The minimum nucleotide distance between candidate donor and acceptor sites are considered likely PEVK exons. We confirmed that each non-overlapping exon was represented by a single reading frame and combined all resultant exons into a single, large exon set. The workflow of PEVK_Finder is summarized in Figure S13. The PEVK_Finder program was written in Python using the Biopython bioinformatics tools package (Cock et al, 2009).

We optimized PEVK_Finder on the well-annotated human and mouse TTN sequences, before applying the tool to a diverse set of mammalian species (Table S1). To determine an optimal set of baseline parameters for PEVK _Finder, we subjected the untranslated DNA reference sequence of human and mouse TTN to a range of parameter settings in PEVK_Finder and compared those results to cDNA-annotated PEVK regions. The complete human and mouse TTN reference DNA were downloaded from the NCBI RefSeq database (O’Leary et al, 2016). The coordinates and DNA sequences for cDNA-annotated human and mouse PEVK exons were gathered from the Ensembl genome browser (Yates et al, 2016; Table S1). Human TTN exons 112-224 were defined as the human PEVK region, as determined by Granzier et al. (2007). Mouse TTN exons 109-207 were defined as the mouse PEVK region, as determined by manual inspection of cDNA-annotated sequences. All possible combinations of minimum exon length (10-30), minimum PEVK ratio (0.45-0.83), and sliding window length (10-30) were used to generate ~17,000 PEVK exon sets per species. The parameter ranges used for optimization testing were determined by examining the minimum and mean exon lengths and PEVK ratios of all human and mouse PEVK exons. To determine the lower boundary for each parameter range, a number slightly below the minimum was chosen, and a number higher than the mean was chosen to determine the upper boundary. Each resulting exon set was compared with the corresponding cDNA annotated exon set for each species using the Basic Local Alignment Search Tool (BLAST; Altschul et al, 1990). Only exons with 100% identity as determined by BLAST were retained for the next steps of optimization. If multiple hits per exon met these criteria, only the hit with the highest bit score was retained.

A match score was calculated to determine how well parameter sets recover the annotated PEVK regions for the ~17,000 parameter combinations per species. The match score is a weighted score that prioritizes exons that recapitulate annotated exons (“recovered exons”; 70%), rewards identical exons (“perfect exons”; 10%), and minimizes exons identified by PEVK_Finder that are not found in the annotations (“extraneous exons”; 20%). *Recovered exons* were calculated as the proportion of the cDNA annotated exons identified by PEVK_Finder and included exons that generated both partial and full BLAST hits; *perfect exons* were defined as the proportion of recovered exons that were 100% identical for the entire length of the annotated exons; *extraneous exons* were calculated by subtracting the number of PEVK_Finder exons with no matches in the annotated database from 100 and dividing that number by 100. When the number of matchless exons found by PEVK_Finder was >100, the extraneous exons score was 0. The final match score was the sum of the three separate weighted scores and ranged from 0 to 1. Parameter space that encompassed the highest match scores were used to determine an optimal set of parameter ranges that yielded the most accurate sets of PEVK exons for a range of mammalian species. All match score scripts were written in R (R Core Team, 2016).

### PEVK Finder validation

To evaluate the utility of PEVK_Finder, we compared the performance of PEVK_Finder on human and mouse TTN with a suite of popular gene annotation tools: Augustus (Stanke & Morgenstern, 2005), FGENESH (Salamov & Solovyev, 2000), geneid (Parra, Blanco & Guigó, 2000) and GENSCAN (Burge & Karlin, 1997). We used each tool with default parameters to predict the exon-intron coordinates of PEVK exons in both the human and mouse TTN sequences. Exon coordinates generated by each tool were manually compared with cDNA-annotated PEVK exon coordinates for the corresponding species. Exons generated by any of the four tools with a partial match (>= 50%) to a corresponding cDNA exon were considered a match, and a tool exon that spanned two or more cDNA exons was considered a single match. cDNA exons with no overlapping coordinates in the tool exon set were considered missing exons. Tool exons with no overlapping coordinates in the cDNA exon set were considered novel exons.

### Phylogeny construction and PEVK region characterization across mammals

We downloaded 43 mammalian TTN sequences representing 16 major orders from the NCBI RefSeq database (Table S1). Our taxonomic sampling is a representative subset from Zhao et al. (2009), which generated a phylogeny of 59 mammals that evolved to fill diverse niches including subterranean, aquatic and nocturnal niches. The genomic TTN sequences for all 43 mammals in the species list were run through PEVK_Finder and used to create novel PEVK exon sets. 41 of these exon sets were used for further analysis. To restrict PEVK exons to the PEVK region, terminal exons were manually determined based on the typical exon proximity and molecular structure patterns observed in the terminal exons of the cDNA-annotated human and mouse PEVK regions. Specifically, the first PEVK exon (human exon 112) has an amino acid sequence similar to “EIPPVVAPPIPLLLPTPEEKKPPPKRI” and is < 3,000 nucleotides before human TTN exon 114, which has an ortholog in all but one species. The last PEVK exon has an amino acid sequence identical or nearly identical to “AKAPKEEAAKPKGPI” and is generally < 3,000 nucleotides after the second to last exon.

Using a full time-calibrated phylogeny of extant mammalian species, we created a phylogeny of the 41 mammalian species in our study to compare PEVK exons across vertebrates and set duck-billed platypus as the outgroup (Kumar et al, 2017). Phylogeny importation and editing was performed in R using the Phytools package (Revell 2012; R Version 3.3.2, R Core Team 2016). To facilitate visual comparisons of PEVK regions, we generated exon-intron diagrams for each species with a custom R script that plotted the coordinates of PEVK exons in each exon set, color-coded by percent PEVK per exon.

PEVK_Finder exons were compared with cDNA-based annotations or *in silico* predicted exons for the 41 mammalian exon sets. The NCBI gene prediction program Gnomon (Souvorov et al, 2010) predicts the optimal coding sequence using a combination of partial alignments and *ab initio* modeling. PEVK_Finder annotations were compared with cDNA-based Gnomon predictions when cDNA was available and were compared with *in silico* Gnomon predictions when no cDNA was available. For simplicity, both sets of predictions are referred to here as “Gnomon exons.” We calculated exons identified by PEVK_Finder or Gnomon, exons that were identified by both (“consensus exons”), and the total number of unique exons identified by both tools. Mean differences between Gnomon and PEVK_Finder were determined using a paired t-test, pairing by species.

### Evolutionary analyses

To facilitate comparisons among PEVK exons, we attempted to identify “orthologous exons”. We use the term “orthologous” to distinguish exons that descend from a common DNA ancestral sequence from those that are related by duplication (i.e. paralogous). Orthologous PEVK exons among species were determined using the reciprocal best BLAST method, with the exception that orthologs were determined for each exon rather than the entire TTN gene (Tatusov, 1997; Bork & Koonin, 1998; Koonin, 2005). The nucleotide sequence of the human PEVK exon set was used as the query and compared with exon sets from the other species. Potential orthologs were called when a given human exon’s top BLASTn species hit (minimum e-value, maximum bit score among hits) returned the original query exon when performed reciprocally. Only orthologs found in the PEVK exon sets of >10 species (including human) were included in further analyses. The remaining sets of orthologous pairs are referred to as “confident orthologs”. Confident orthologs were aligned by codon with the ClustalW (Goujon et al, 2010) algorithm as implemented in MEGA7 (Kumar et al., 2016), using all other default parameter settings. Codons missing >15% of data among species were eliminated from the alignment. We estimated average evolutionary divergence over all sequence pairs for each confident ortholog following Tamura et al., (2004) as implemented in MEGA7 (Kumar et al., 2016). Values for each ortholog were calculated in MEGA7 with default parameters and 100 bootstrap replications.

We also evaluated repetition and duplication within the PEVK region by comparing all PEVK exons for a single species with each other. For a given species, all possible exon pairs in the PEVK exon set were aligned and pairwise substitutions per exon were calculated using MEGA Proto and MEGA-CC (Kumar et al, 2012). The PEVK region was divided into the subregions PEVK-N and PEVK-C, defined in *Results: PEVK region hierarchical structure*. The pairwise exon alignments were divided into quadrants representing PEVK-C:PEVK-N (Quadrant I), PEVK-C:PEVK-C (Quadrant II), PEVK-N:PEVK-N (Quadrant III), and PEVK-N:PEVK-C (Quadrant IV) comparisons. For each species, we calculated mean substitutions per exon for all quadrants and results were compared with a one-way ANOVA and a Tukey’s HSD post-hoc test. To evaluate repetition at the nucleotide level, we concatenated exonic sequences from the human PEVK-C region and generated a dot plot in the Dotlet web browser (Junier and Pagni, M., 2000).

We tested for positive selection acting on individual codons using site models as implemented in the Phylogenetic Analysis by Maximum Likelihood (PAML) software package (Yang, 2007). For each ortholog alignment, a phylogenetic tree was estimated using the M0 model, then models M0 (one ratio), M1a (neutral), and M2a (positive selection), were run. For each model, ω was set first at 0.5, then at 1 and then lastly at 3. We recorded Lnl values for each model and test statistics for two model comparisons: M0 vs. M1a and M1a vs. M2a. The PAML χ2 calculator was used to determine the p-value of each comparison. All sites under selection, as determined by Bayes Empirical Bayes analysis (Yang et al., 2005), were recorded for estimates with posterior probabilities > 0.5.

Finally, we tested for charge shifts in codons under positive selection. For each codon, the corresponding amino acid for each species was assigned one of four values based on residue charge: hydrophobic/uncharged, polar, positively charged, or negatively charged. The ancestral charge status of each node in the corresponding mammal tree was estimated using an equal rates (ER) model. Ancestral state reconstruction was performed in R using the Phytools package (Revell 2012; R Core Team 2016).

### Data Availability

The PEVK_Finder code is available for download at https://github.com/kmuenzen/pevk_finder_public. Tables S1 and S2 contain names and NCBI accession numbers for sequences used in this study.

## RESULTS

### PEVK_Finder parameter optimization

PEVK_Finder was developed to annotate the PEVK region of TTN across mammals. We determined optimal parameters (exon length, window size, and PEVK ratio) for downstream applications by comparing PEVK_Finder results with the human and mouse annotations and calculating a match score. For each human and mouse model, we tested a total of ~17,000 parameter combinations. Optimal match scores for human were achieved when minimum exon length was 10-15 nucleotides, PEVK ratio was 0.54-0.55, and window length was 10 nucleotides (Figure 1a). Optimal match scores for mouse were achieved when minimum exon length was 12-14 nucleotides, PEVK ratio was 0.53-0.54, and window length was 10 nucleotides (Figure S1a). Consensus parameters were therefore set as follows: window length = 10 nucleotides, PEVK ratio = 0.54, and minimum exon length = 12 nucleotides.

**Figure 1.**
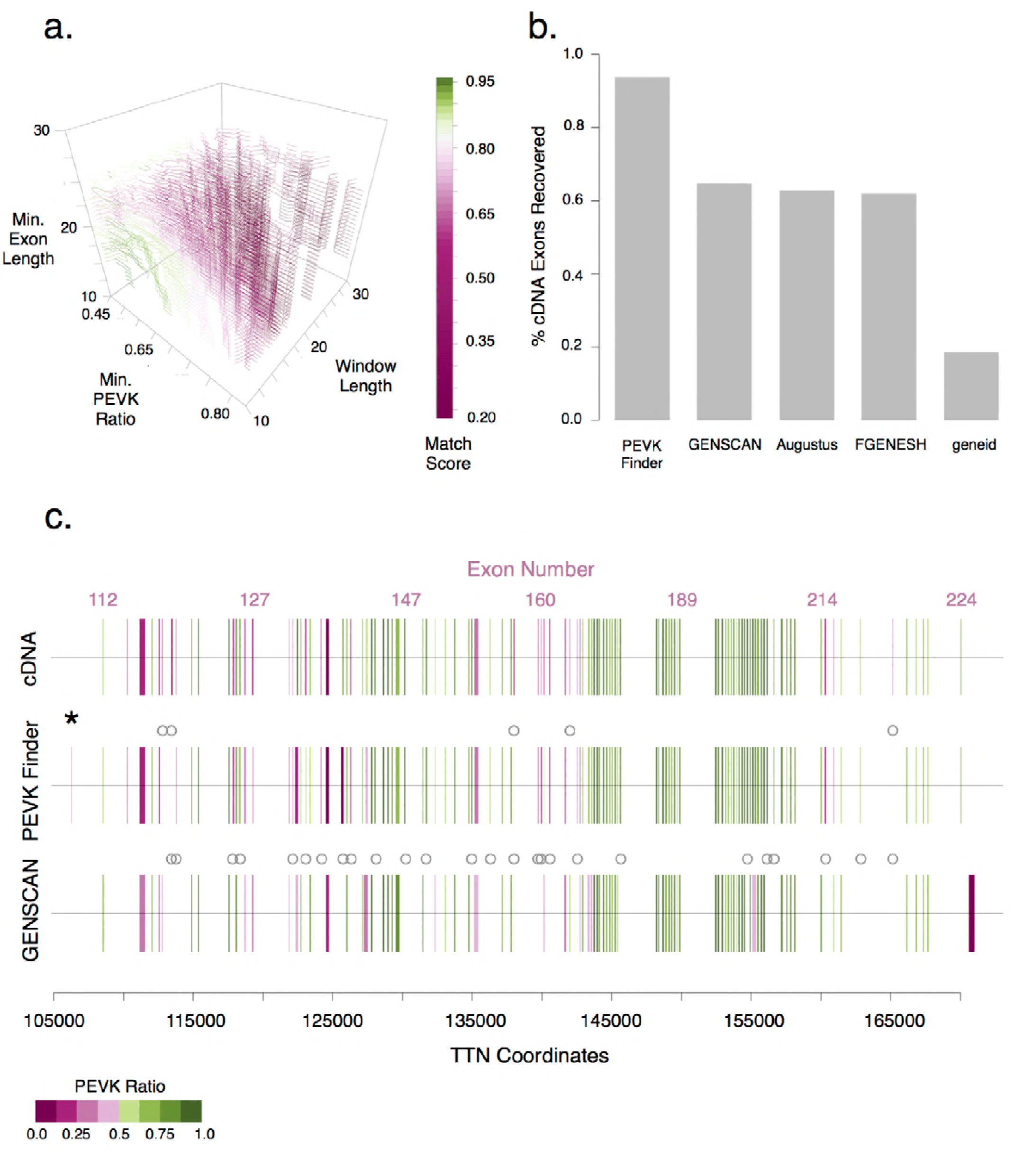
PEVK Finder tool optimization and evaluation of human TTN. **a)** Match scores of human PEVK exon sets were generated using different combinations of minimum exon length, PEVK ratio and sliding window length ratio parameter settings. **b)** PEVK_Finder recovered more PEVK exons than other existing gene prediction tools (GENSCAN, Augustus, FGENESH and geneid). **c)** PEVK_Finder outperformed GENSCAN at recovering the exon-intron distribution of human TTN PEVK exons identified by cDNA. Lines indicate exon boundaries and the thickness of the lines is determined by the exon coordinates. Grey circles indicate exons that were missed by either PEVK Finder or GENSCAN, and black asterisks indicate automatically annotated exons that were not annotated by cDNA. The PEVK ratio scale is giving in the figure.

Using the optimal parameter values, PEVK_Finder identified 109 exons in the PEVK region in human TTN (113 annotated) and 93 PEVK exons in mouse TTN (99 annotated). In humans, 106 PEVK_Finder exons recovered annotated exons (94%) and 70 of these were perfect matches. Of the 36 non-perfect exon matches, 7 were overestimations of exon length and 29 were underestimations. When PEVK_Finder exon length was >50% reduced relative to the annotated length, the annotated exons were often long, with low PEVK ratios. In these cases, PEVK_Finder tended to truncate the exons in regions due to low PEVK ratio (e.g., human TTN exon 114). When PEVK_Finder overestimated exon length, all PEVK_Finder exons were <100 nucleotides long and length differences were marginal (3-4 amino acids difference). Patterns were similar with the mouse TTN, with the exception that overestimations were more frequent (12 of 37 non-perfect exon matches) due to regions of low PEVK ratios (Fig S1c).

### PEVK_Finder validation

To evaluate the utility of PEVK_Finder, we compared results generated from PEVK_Finder with four different gene annotation tools. Overall, PEVK_Finder outperformed all other tools when identifying PEVK exons in the TTN gene. PEVK_Finder identified 97% of human TTN exons, while no other tool identified more than 81% of human PEVK exons (Fig 1b). PEVK_Finder also missed far fewer exons than GENSCAN in both human and mouse TTN (grey circles Figure 1c, S1c). PEVK_Finder was also the only tool that identified >80% of both human and mouse exons in the PEVK region (Figures 1b, S1b). FGENESH also identified a high proportion of mouse exons (81%), but was far less effective than PEVK_Finder at annotating human exons (62%). PEVK_Finder performed slightly better on human TTN (94%), than mouse TTN (90%). PEVK_Finder also annotated exons across a range of PEVK ratios and in regions of short, high-density exons (Figures 1c, S1c); these regions were a challenge for GENSCAN. In the few instances when PEVK_Finder did fail to annotate exons, it was usually those exons with low PEVK ratios. GENSCAN also tended to lump distinct exons together, likely due to missed alternative splicing sites. PEVK_Finder, FGENESH, and GENSCAN could be applied to both mouse and human TTN, whereas geneid and Augustus are only suitable for human data. In addition to identifying known exons, PEVK_Finder discovered novel putative PEVK exons in both the human (1; Figure 1c) and mouse (2; Figure S1c) TTN sequences.

We extended our evaluation of PEVK_Finder by evaluating its performance on other species with annotated TTN sequences. Of the 41 mammal species tested, only five have cDNA data for TTN in NCBI (Table S2). Human TTN has the most complete annotation, with consensus cDNA from 11 isoforms. The American Pika (*Ochotona princeps*) has cDNA data from 5 isoforms, the house mouse (*Mus musculus*) and the Gairdner’s Shrewmouse (*Mus pahari*) have cDNA data from 3 isoforms each, and the Chinese Rufous Horseshoe Bat (*Rhinolophus sinicus*) has cDNA data from 1 short isoform. For the remaining 36 species, we compared PEVK_Finder results with the annotations provided by NCBI generated by the Gnomon program (Souvorov et al, 2010). Across all species, PEVK_Finder identified significantly more exons (mean: 89.4, SD: 8.33) than the Gnomon-based annotations (mean: 73.9, SD: 10.6; t = 9.37, df = 40, *P* = 1.23*10-11; Table S2). The number of exons identified by PEVK_Finder represents a significantly higher percentage of the total exons (mean: 93.6%, SD: 2.35) than Gnomon-based annotations (mean: 76.7%, SD: 9.29; *t* = 11.10, *P* = 8.71*10-14; Table S2). PEVK_Finder also identified many more putative novel PEVK exons that were not described previously by either Gnomon or cDNA (mean: 22.7; SD: 10.17).

### PEVK region hierarchical structure

PEVK_Finder successfully generated exon sets for 41 of the 43 species tested; we were unable to obtain exon sets for two species (*Sorex araneus* and *Leptonychotes weddellii*) due to non-canonical splicing sequences that were not compatible with those used by PEVK Finder to identify exon-intron boundaries. The PEVK region of the remaining 41 species exhibited structural similarities in both exon-intron spacing and PEVK content per exon (Figure 2). The total number of exons identified by PEVK_Finder ranged from 81 – 116 exons, 74 – 109 of which were within the bounds of the exons defining the start and end of the PEVK region (Table S2). Interestingly, the three species with the smallest number of PEVK exons are all aquatic diving mammals (*Odobbenusn rosmarus –* walrus (82), *Tursiops truncates –* dolphin (81), *Neomonachus schaunslandi* –Hawaiian monk seal (82)). However, two of these species also show gaps in the PEVK regions, which may be real or may be due to assembly artifacts (see below).

**Figure 2.**
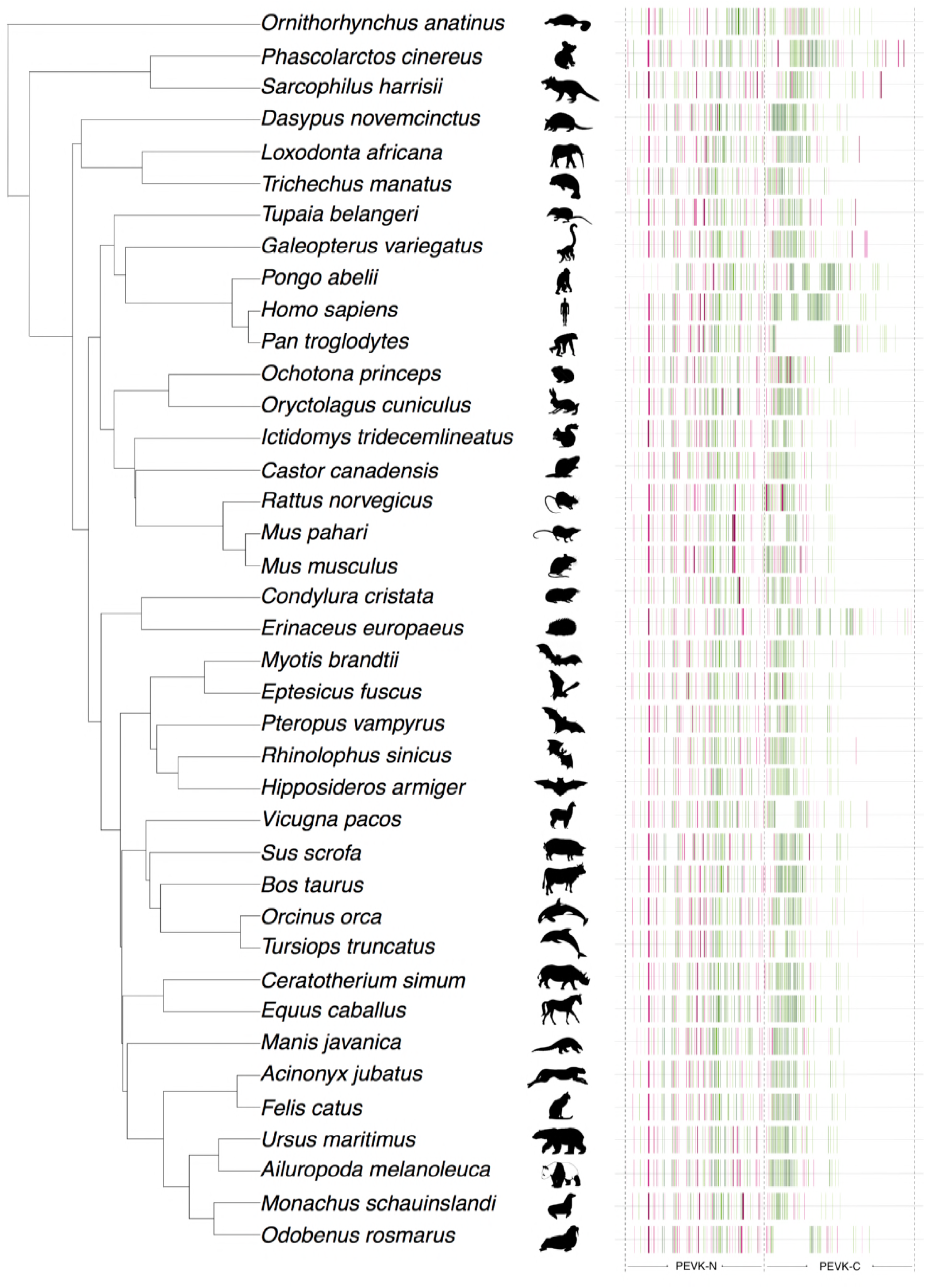
Phylogenetic comparison of PEVK exon structure. PEVK_Finder exon-intron plots were overlaid onto a time-calibrated phylogeny for 39 mammalian species. The dotted line indicates the approximate boundary of PEVK-N and PEVK-C segments of the PEVK region, based on the human boundary between the two segments. PECK ratio scale is the same as in Fig. 1.

Exon structure of the upstream ~half of the PEVK region is relatively conserved over mammalian evolution, while the downstream ~half varies considerably. Based on this observation, we divided the PEVK region into two distinct regions: a structurally conserved region (“PEVK-N”; human exons 112– 165) and a structurally variable region (“PEVK-C”; human exons 166 – 224). Because the end of the PEVK-N region and the beginning of the PEVK-C region varies 1-5 exons between species, a visual landmark was used to manually determine the PEVK-N/C boundary for each species. The manually determined PEVK-N/C boundary is preceded by 3-4 very closely spaced, low PEVK-ratio exons, and is followed by 3-4 more low-PEVK ratio exons and a dense, high PEVK-ratio region (Figure 2). Across mammals, intron/exon boundaries and the number of exons are more similar across the PEVK-N (mean: 50.00, SD: 2.51) versus the PEVK-C region (mean: 39.46, SD: 1.27, *f* = 0.10, *P* < 1.70E-11) (Figure 2, Table S2). Several species (e.g., chimpanzee, dolphin, walrus and alpaca) also show large deletions of PEVK exons in the PEVK-C region. The walrus TTN sequence contains a large string of >9,000 ‘N’ bases, indicating that the gap in the walrus PEVK-C region is likely due to low assembly quality in this region. Alpaca TTN also contains several strings of N’s that range from 500-5,000 bases long, which are likely responsible for the large gaps. However, while several small strings of 10-150 ‘N’ bases are found near the PEVK-C region of chimpanzee and dolphin TTN, no large strings of Ns are present. The large deletions in the chimpanzee and dolphin may therefore represent true PEVK exon “deserts”. Alternatively, the gaps could also be an artifact of non-canonical splice sites or small strings of N’s interfering with splice site detection or PEVK ratios. Complete genomic sequences of TTN and/or RNA-based annotations could improve resolution of PEVK exons in these species.

### PEVK region sequence variability within an individual

Although the PEVK-C region is structurally variable across mammals, its sequence content is more similar within a given individual than the PEVK-N region (Figures 2,3). Exons in the PEVK-C region tend to be short (mean: 75.76, SE: 0.685) with high PEVK ratios (mean: 0.84, SE: 0.003), whereas those in the PEVK-N region consist of a range of lengths (mean: 93.29, SE: 1.37: *t* = 11.44, *f* = 8.68, *P* < 2.2*10-16) and lower PEVK ratios (mean: 0.79, SE: 0.003: *t* = 12.84, *P* < 2.2*10^-16^) (Figure 2). Likewise, exon-exon identity comparisons (Figure 3a) and dot plots of nucleotide sequence (Figure 3b) reveal large blocks of highly repetitive sequence in the PEVK-C region. When looking across all species, the mean substitutions per exon varied significantly among the four quadrants (one-way ANOVA: *F*3,160 = 33.68; *P* = 2 * 10-16; Figure 3c). Post hoc comparisons using the Tukey HSD test indicated that the mean number of substitutions per exon for PEVK-C:PEVK-C exon comparisons (mean 1.02; SD: 0.17; Quadrant II) were significantly lower than comparisons for PEVK-C:PEVK-N exons (mean 1.26; SD: 0.12; *P* < 0.0001; Quadrant I), PEVK-N:PEVK-N exons (mean: 1.23; SD: 0.10; *P* < 0.0001; Quadrant III), and PEVK-N:PEVK-C exons (mean 1.26; SD: 0.12; *P* < 0.0001; Quadrant IV). No other quadrant:quadrant comparisons were significantly different (*P* > 0.05 in all cases). Together, these analyses suggest that exons within the PEVK-C region have more similar sequences, while exons in the PEVK-N region are more variable in sequence.

**Figure 3.**
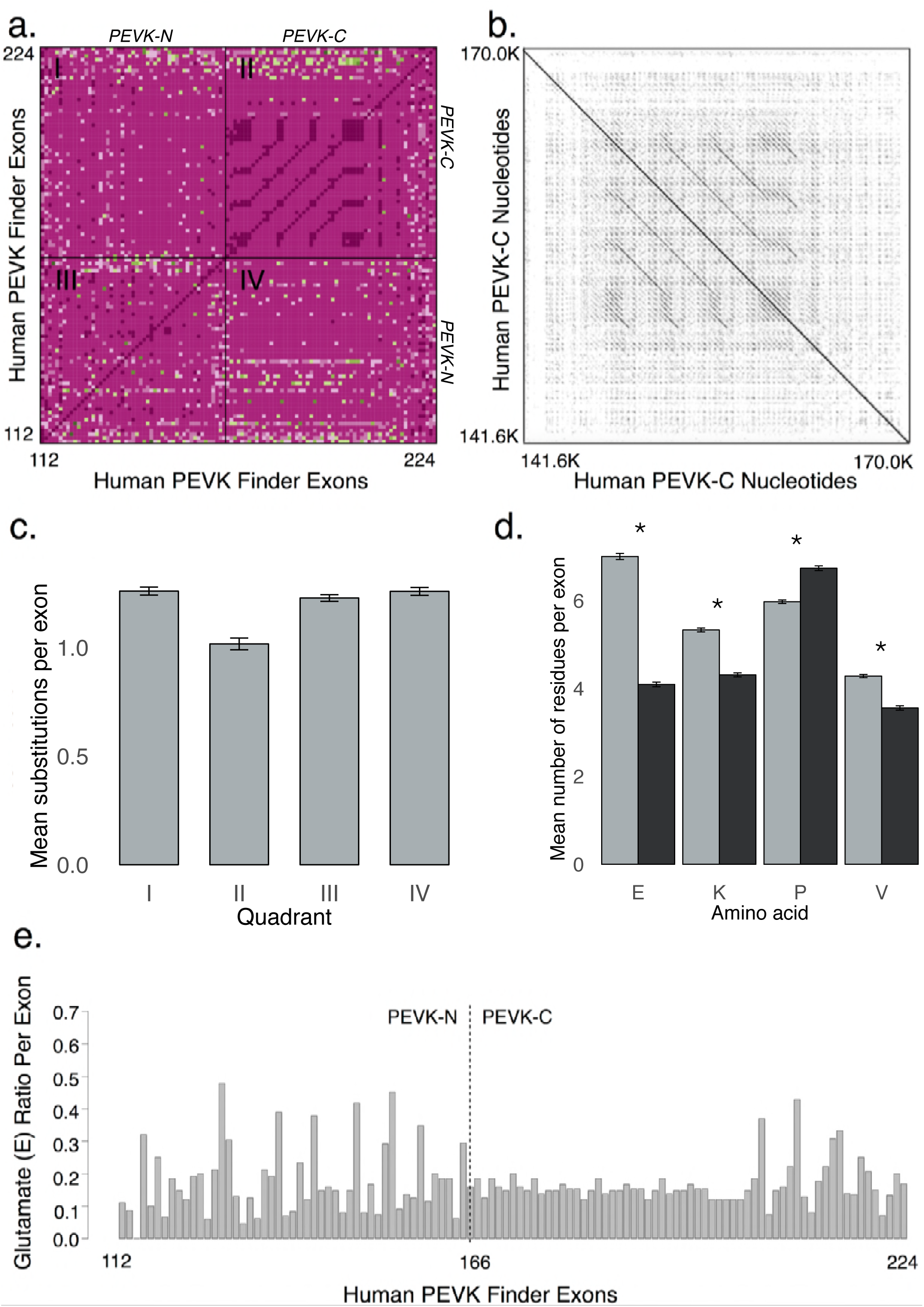
Exon and nucleotide-level comparison of PEVK-N and PEVK-C regions. **a)** A heat map of substitutions among PEVK exons within a species: I) PEVK-N vs. PEVK-C, II) PEVK-C vs. PEVK-C, III) PEVK-N vs. PEVK-N, and IV) PEVK-C vs. PEVK-N. Dark pink indicates exon pairs with few substitutions, whereas light pink and green indicate exons pairs with many substitutions. In general, PEVK-C exons are highly repetitive and homogeneous, while PEVK-N exons are more variable**. b)** A reciprocal dot plot of the human PEVK-C nucleotide sequence shows the repetitive nature of the PEVK-C region in humans (Dotlet, https://dotlet.vital-it.ch). **c)** Mean pairwise substitutions per exon across all 39 species for each quadrant from Figure 3a. Bars represent mean ± s.e. **d)** Mean P,E,V,K amino acids per exon across all 39 mammalian species. There is significantly more glutamate (E), valine (V) and lysine (K) per exon in the PEVK-N region and more proline (P) per exon in the PEVK-C region across all 41 species. Grey bars: PEVK-C exons; Black bars: PEVK-N exons. Bars represent mean ±s.e. Asterisks denote significance at P < 0.05. e) Ratio of glutamate (E) per exon in human TTN PEVK. There is relatively more glutamate (E) per exon in the PEVK-N region compared to the PEVK-C region. The dotted line indicates the boundary between the PEVK-N and PEVK-C regions.

The PEVK-N and PEVK-C regions also encode for significantly different amounts of the core P, E, V, and K amino acids. PEVK-N exons contain significantly more glutamate (1.7X; *t* = 33.51, *P* < 6.84E-31), lysine (1.2X; *t* = 15.76, *P* < 9.32E-19), and valine (1.2X; *t* = 13.81, *P* < = 8.30*10-17) amino acids than PEVK-C exons (Figure 3d). PEVK-C exons contain significantly more proline amino acids (1.1X; *t* = 13.83, *P* < 7.79*10-17). Non-PEVK amino acid residues, such as isoleucine, arginine, threonine, leucine, tyrosine and serine, were also more common in PEVK-N exons (Figure S4).

### Evolutionary analysis of the PEVK region across mammals

We performed a reciprocal best BLAST (Tatusov 1997; Bork and Koonin 1998; Koonin 2005) to detect orthologs in the PEVK region. Overall, orthology across the PEVK region is low, with only 22 confident orthologs detected in the PEVK-N region (43% of human PEVK-N exons) and 13 in the PEVK-C region (22% of PEVK-C exons) (Figures 4, S3). More orthologs are detected in the PEVK-N region than the PEVK-C region (red/orange, Figure S3), consistent with a more conserved exon structure in the PEVK-N region. Exons in the PEVK-C region display greater sequence divergence across the mammalian tree (mean: 0.11; SE: 0.01) than the PEVK-N region (mean 0.08; SE: 0.01), though this difference is not significant (*t* = 1.67, *P* = 0.11) and primarily driven by three exons in the PEVK-C region (Figure S3).

**Figure 4.**
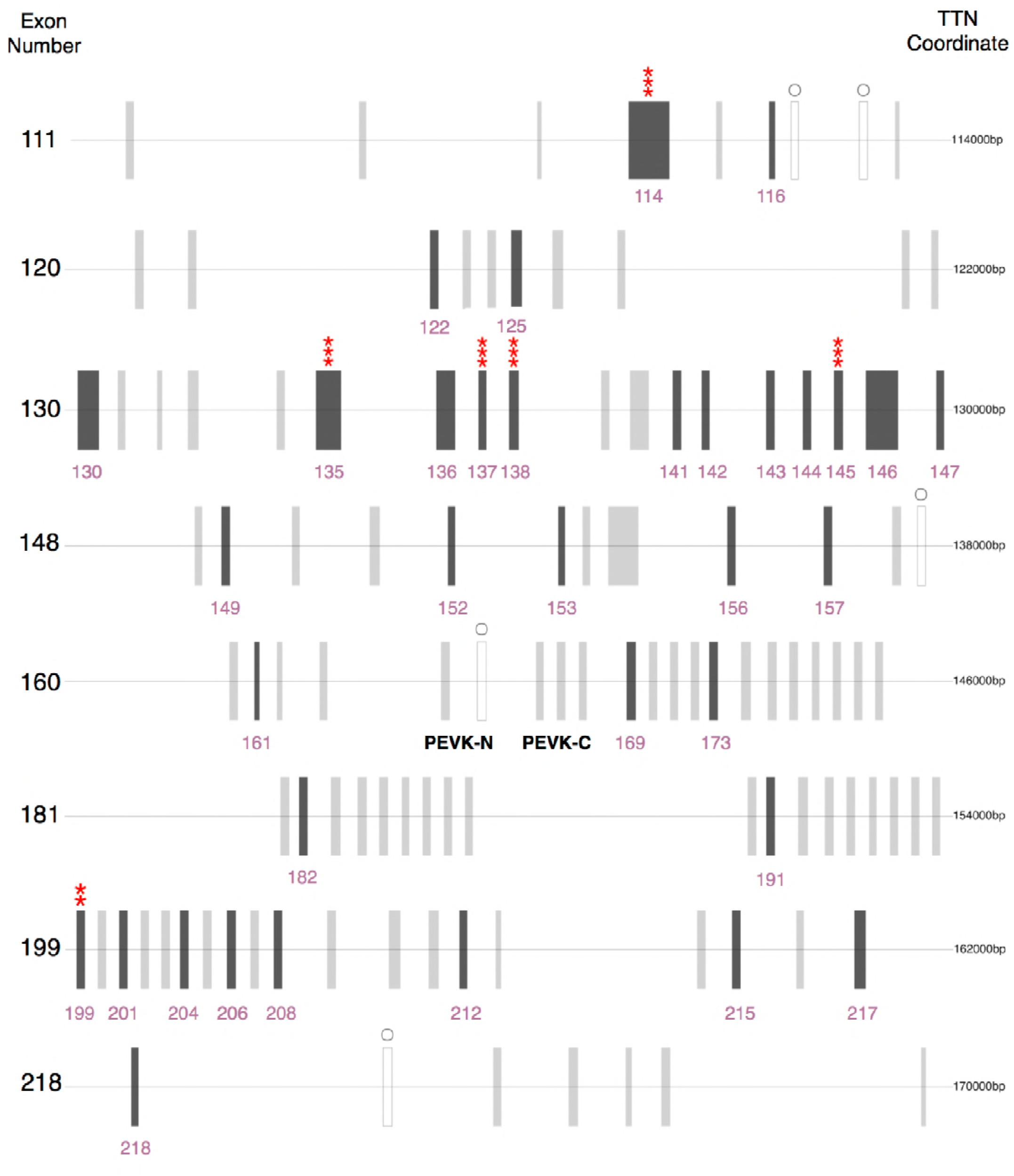
Human PEVK_Finder exon-intron plot depicting orthologous exons and codons under significant positive selection. Dark grey bars (labeled with their respective exon numbers) indicate orthologous exons, grey circles indicate exons missed by PEVK Finder, and asterisks indicate codons under selection at varying levels of significance. Asterisks indicate significance; * P > 0.5; ** P > 0.75; *** P > 0.95. Only the most significant codon for a exon is noted. Exons 114, 116, 122, 125, 130, 135, 136, 137, 138, 141, 142, 143, 144, 146, 152 153, and 161 are either constitutively expressed or expressed in >95% of TTN transcripts in skeletal muscle according to Savarese et al. 2018. Details about codons under selection are in Figures S5-S12).

We also examined the nature of selection acting on codons in the PEVK region. Specifically, we tested for positive selection using random-sites models in PAML, which allowed ω to vary among sites but not lineages. For six sets of orthologous exons (human exons 114, 135, 137, 138, 145 and 199), models that allowed certain codons to evolve under positive selection (ω > 1; M2a) were significantly better than models that restricted codons to be conserved (0 < ω < 1) or neutral (ω = 1; M1a; Figure 4, Table 1). Model M1a vs. M0 was also significant for six exons investigated (results not shown). Five exons with sites under positive selection were in the PEVK-N region, and one was in the PEVK-C region. In total, 20 codons were under positive selection (Figures S5-S10; Table 1). Results were qualitatively similar for all initial ω values.

The polarity of amino acids can influence protein-binding interactions and solubility, and therefore may contribute to adaptive evolution of titin. Therefore, we used ancestral state reconstruction to examine charge transitions in TTN exons with confident orthologs. Our analyses show that codons 42 of exon 114 and 6 of exon 135 are under positive selection and have substitutions that change the charge of the codon (Figures S11-S12). Codon 42 of exon 114 features shifts from hydrophobic to polar amino acids from a minimum of 3 to ~6 times, with clear independent shifts occurring in cetaceans, bats, and beaver. This codon also shifts from a hydrophobic to a positively charged amino acid in *Mus musculus*. Codon 6 of exon 135 reveals four independent transitions from positively charged to polar amino acids across mammals. Intriguingly, in cetaceans, this codon shifts from positively charged to hydrophobic. Thus, in both instances of codons with evidence for charge transitions, cetaceans are one of the few groups to experience changes in polarity and sometimes do so in unique ways (Figure S12). Together, these charge changes and low PEVK exon numbers suggest that titin in cetaceans and other aquatic diving mammals may be under different selective pressures than terrestrial mammals.

## DISCUSSION

### PEVK_Finder improves the annotation of the PEVK region across mammals

The PEVK region of TTN possesses many attributes that confound standard gene prediction tools; it contains numerous short, repetitive exons that vary within and across species in sequence content, exon structure, and length. However, it is this very complexity that makes the PEVK region a promising source of raw material for evolutionary adaptation of muscle (Freiburg et al. 2000; Tompa 2003; Prado et al. 2005). Robust annotations of the PEVK region are therefore needed to link molecular evolution of TTN with physiological performance over evolutionary time.

While accurate gene prediction may be achieved with RNA-Seq or similar methods, relying on transcriptomic data for gene prediction is not always feasible due to time and cost constraints. For example, only 5 of the 43 species examined in this study had cDNA data supporting their TTN annotations at the time of the study. The plethora of alternatively spliced isoforms of TTN further complicates RNA-based gene predictions, as data from numerous tissues, developmental stages, and/or individuals are required to generate complete annotations (e.g., Savarese et al 2018). To this end, we developed PEVK_Finder, which combines *ab initio* and signature-based approaches to gene prediction and targets the PEVK region of TTN (Wang et al 2004; Lerat 2010; Sleater 2010). Our tool depends solely on nucleotide sequence and searches for exons with motifs and patterns common to well-annotated PEVK regions (i.e., human, mouse). Similar custom tools have been developed for classes of transposable elements, which are also highly repetitive and often missed by standard annotation tools (Lerat 2010).

Here, we find that PEVK_Finder consistently improves the annotation of the PEVK region across mammals. One major challenge of annotating repetitive regions with short exons is that exons are commonly missed. PEVK_Finder consistently detects significantly more PEVK exons than standard annotation tools, both in model and non-model species (Figure 1b, 1c; Table S2). Specifically, PEVK_Finder outperforms other tools in regions of short, high-density exons and across broad ranges of PEVK ratios. PEVK_Finder also identifies numerous putative PEVK exons that are not found in cDNA annotations or identified by other bioinformatics tools (Figure 1c; Table S2). It is possible that these novel exons are not transcribed, but TTN has many alternatively spliced isoforms that may not be completely characterized by cDNA. For example, manual annotation of genomic TTN sequences has discovered novel TTN exons in the past (e.g., Freiburg et al. 2000; Granzier et al., 2007).

While PEVK_Finder improves available annotations in non-model species, we caution that it does not perfectly replicate the PEVK region in human and mouse TTN. PEVK_Finder occasionally misses, truncates, or joins distinct exons, albeit at a lower frequency than the other tools tested. Exons with low PEVK ratios, internal splice sites and non-canonical splice sites cause most of these errors. Indeed, we were unable to obtain exon sets for two of the 43 species we tested due to the presence of non-canonical splice sites. In the future, incorporation of hidden Markov models into the tool could improve issues with splicing and variable PEVK ratios. Given this, we argue that PEVK_Finder is most useful for determining major trends in PEVK structure and evolution in non-model species, as exemplified here. When complete, perfectly resolved annotations are required in a non-model species, PEVK_Finder should be used in conjunction with RNA-Seq or cDNA sequencing from numerous tissues, individuals, and developmental times. Likewise, pairing PEVK_Finder with another annotation tool, such as Gnomon, could also provide more comprehensive annotations (Table S2).

### The evolution of the PEVK region

Our results reveal contrasting patterns of constraint and divergence across the PEVK region, suggesting two subregions with distinct evolutionary dynamics. The PEVK-N region shows relatively conserved length and exon structure over evolutionary time, but evidence of diversifying selection and more variable amino acid content (Figures 2, 4). In contrast, the PEVK-C region varies dramatically in length and exon number across mammals, but these exons tend to be more similar (Figure 2, 3a, 3c). Overall, the documented patterns argue for selection maintaining particular, “essential” PEVK-N exons over evolutionary time, with diversifying selection targeting specific codons. For example, 17 out of 22 PEVK-N exons with confident orthologs are either constitutively expressed or expressed in >95% of TTN transcripts in skeletal muscle (Figure 4).

Conservation over both evolutionary time and across isoforms suggest these exons may play key roles in vertebrate muscle function. Additionally, five PEVK-N exons have codons that experienced adaptive evolution and may be implicated in diversification of muscle function over mammalian evolution. Specifically, codons in exons 114 and 135 experienced major shifts in charge over mammalian evolution and may influence titin-protein binding. Conversely, expansion and contraction of the total length of the PEVK-C region, rather than selection on any particular exon, dominates the evolution of the PEVK-C. Such length variation has the potential to contribute to functional differences, as altering the size of the titin “spring” through alternative splicing of more or fewer PEVK repeats can affect titin’s compliance, though neutral processes may also be relevant (reviewed in Linke 2018). Future work can focus on disentangling the effects of natural selection acting on specific codons from PEVK length variation, and how both contribute to evolutionary adaptation of TTN.

What explains the discrepancy between the molecular patterns documented in the PEVK-N and PEVK-C regions? Intraspecific sequence comparisons of PEVK exons offer clues. Within human titin, the sequence content of the PEVK-C is more repetitive than the PEVK-N (Figures 3a, 3b, 3c, S5). This suggests a mechanistic basis for expansion and contraction of the PEVK-C over the course of mammalian evolution. Replication slippage and recombinatorial repeat expansions often occur in regions with tandem repeats, and may result in exon duplication and loss (Tomba 2003). Mobile LINE element insertions have also been implicated in TTN remodeling by causing PEVK exon duplication or differential splicing (Granzier et al. 2007). Because we find that the PEVK-C contains more repeats than the PEVK-N, we argue that any process generating exon duplication or loss occurs more frequently in the PEVK-C.

### Implications for titin function

The length and exon structure of the PEVK region varies remarkably over evolutionary time, with implications for titin functionality. The PEVK region of titin has long been known to contribute to passive stiffness of muscle (Gautel and Goulding, 1996, Linke et al., 1998). Through alternative splicing, titin can be expressed as isoforms with varying lengths of the PEVK domain which correlate with the passive properties of different muscle types (Freiburg et al., 2000, Prado et al., 2005). To date, at least twelve titin isoforms have been identified in cardiac and skeletal muscles from human, mouse, and rabbit tissues. Muscles with long PEVK segments have low passive stiffness whereas muscles with shorter segments have higher passive stiffness (Freiburg et al., 2000, Prado et al., 2005). The repetitive nature of PEVK-C exons may allow for variability in the length and stiffness of titin isoforms. Repeats within the PEVK region have been postulated to be functionally equivalent to fulfill entropic chain requirements (Tomba 2003). While this may be the case for repeats within the PEVK-C region, it is less clear for repeats within the PEVK-N region. Alternatively, variation in exon number across the PEVK region may be a product of neutral processes that result in exon skipping and loss, without affecting TTN function.

While a mechanism has not yet been elucidated, several researchers have proposed that during muscle activation, titin stiffness increases in the presence of calcium possibly by titin-actin binding (Herzog et al., 2012, Nishikawa et al., 2012, Schappacher-Tilp et al., 2015). Within the PEVK region, exons can be composed of strings of negatively charged glutamate residues (E-rich motifs). Results from single molecule experiments have shown that E-rich PEVK segments bind to actin filaments, which produces a viscous load that could possibly resist the sliding of the thin filament along the thick filament during muscle contraction (Kellermayer and Granzier, 1996, Labeit et al., 2003, Nagy et al., 2004, Bianco et al., 2007). If this interaction is necessary for increasing titin stiffness in activated muscle, splicing of exons that encode for amino acids responsible for changing charge or binding affinity to actin could have dramatic effects on titin function. For example, several studies have implicated titin in the enhancement of active muscle force with stretch (Leonard and Herzog 2010, Powers et al., 2016, Hessel et al., 2017). If titin stiffness failed to increase as a result of a change in amino acid charge and/or binding to actin or other proteins, muscles could show decreased force during stretch and alter an organism’s ability to move. Our data show that PEVK-N exons are conserved and have a higher percentage of E-rich motifs than PEVK-C exons. In addition, PEVK-N exons 114 and 135 exhibit charge shifts that could affect PEVK interactions with other proteins. However, no studies to date have shown calcium dependent PEVK-actin binding (Bianco et al., 2007, Nagy et al., 2004, Linke et al., 2002). Thus, it is unlikely this interaction underlies the increase in titin stiffness during muscle activation. It remains unclear how E-rich motifs or change in charge may affect titin function. Further studies on the expression of PEVK-N exons could serve to elucidate the role of the PEVK in active and passive muscle.

### Conclusions

In summary, PEVK_Finder provides a useful method for determining major trends in PEVK structure and evolution in non-model species. By characterizing the PEVK region of titin in mammals, we identify two potential pathways through which selection could shape titin function—by changing amino acid sequences of specific codons and length variation in the size of the PEVK region. In the PEVK-N, exons with confident orthologs that are constitutively expressed in isoforms are strong candidates for future work testing the essentiality of particular exons underlying titin stiffness and muscle function. Together, the construction of custom annotation tools for challenging-to-annotate genes can facilitate novel insight into the diversification and nature of selection acting on important proteins like titin.

## Acknowledgements

We thank Kiisa Nishikawa, Jocelyn Crawford, Silvia Leblanc, Keon Rabbani and Emma Bekele for their helpful comments on earlier versions of this manuscript. This work was supported by the National Science Foundation (IOS-1731917 awarded to J.A.M.) and start-up funds for F.R.F. and J.A.M.

